# A Bayesian Approach to Correcting the Attenuation Bias of Regression Using Polygenic Risk Score

**DOI:** 10.1101/2023.11.27.568907

**Authors:** Geyu Zhou, Xinyue Qie, Hongyu Zhao

**Author notes:** These authors contributed equally to this work (Geyu Zhou, Xinyue Qie).

## Abstract

Polygenic risk score (PRS) has become increasingly popular for predicting the value of complex traits. In many settings, PRS is used as a covariate in regression analysis to study the association between different phenotypes. However, measurement error in PRS causes attenuation bias in the estimation of regression coefficients. In this paper, we employ a Bayesian approach to accounting for the measurement error of PRS and correcting the attenuation bias in linear and logistic regression. Through simulation, we show that our approach is able to obtain approximately unbiased estimation of coefficients and credible intervals with correct coverage probability. We also empirically compare our Bayesian measurement error model to the conventional regression model by analyzing real traits in the UK Biobank. The results demonstrate the effectiveness of our approach as it significantly reduces the error in coefficient estimates.

## Introduction

Genome-wide association studies (GWAS) have generated a wealth of data over the last two decades (1). A common practice of extracting information from GWAS data is to construct the polygenic risk score (PRS) by aggregating the number of risk alleles weighted by the effect size for each single nucleotide polymorphism (SNP) across the genome. PRS can be viewed as the genetic prediction of complex traits and has great promise in precision medicine for identifying individuals with higher disease risk (2). With the development of large-scale biobanks, another important application of PRS is to explore the relationship between different phenotypes (3, 4). In this setting, because covariates may not be directly observed for some individuals, researchers sometimes first use PRS as the predicted value of the covariate and then perform regression to estimate the coefficient with other observed outcomes. For example, studies have found that PRS of lipid traits are associated with coronary artery diseases (CAD) (5, 6).

PRS is often regarded as a noisy predictor of true phenotypes due to the finite sample size of GWAS and the complexity of genetic architecture. Currently the prediction accuracy of PRS is moderate for most traits, explaining only 5 to 30 percent of phenotypic variance (7, 8). It is well known that measurement error in one or more covariates results in the attenuation bias (i.e. bias towards 0) when estimating the regression coefficients (9). Hence, as we will show, employing PRS as a replacement of the actual covariate may lead to a diminished estimate of the regression coefficient.

The study of measurement error in the regression setting has been an interest to statisticians for many decades (9). In the linear regression setting, the expected value of the least squares estimate subject to the measurement error is the true value multiplied by the reliability ratio, which is defined as the variance of the observed covariate divided by the variance of the true covariate (9). If the reliability ratio is known, a consistent estimator of the coefficient would be the least squares estimate divided by the reliability ratio. The setting of PRS slightly differs from this traditional setting of measurement error model from two aspects. First, the slope of the PRS regressing on the actual phenotype is typically not equal to 1 (7). Second, the logistic regression model may be used when the outcome is binary. Consequently, we took a Bayesian approach instead to handle the measurement error of PRS in the regression model due to its ability to accommodate more flexible assumptions (10).

In this work, we first describe our setting of the Bayesian measurement error model for PRS. We performed simulation to investigate whether our approach is able to reduce the bias in linear and logistic regression. Finally, we apply our approach to eight pairs of observed traits in the UK Biobank (UKBB) to show that our method is able to reduce the attenuation bias.

## Methods

### Outcome Model

We assume that testing data of outcome *Y*_*i*_ are available for *N* individuals, where *i* = 1, …, *N*. For individual *i*, let *Z*_*i*_ denote the covariate of interest subject to measurement error and *W*_*i*_ denote the matrix of error-free covariates (e.g. age and sex). Both outcome and covariates are assumed to be standardized to have mean 0 and variance 1 unless they are binary. We consider two types of outcome models for *Y*_*i*_— linear and logistic regression. For a linear regression model, we have

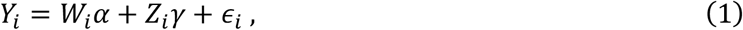

where α is the vector of effect size corresponding to covariates *W*_*i*_, *γ* is the effect size of the covariate of interest, and *ϵ*_*i*_ ∼ *N*(0,σ^5^) is the random error.

Similarly, for a logistic regression model we have

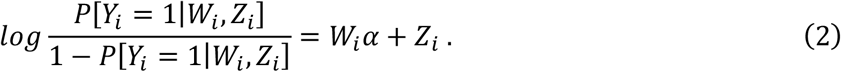

### Measurement Error in PRS

We assume that the covariate of interest *Z*_*i*_ is not observed for *i* = 1, …, *N* individuals. Instead, we have access to *Z*_*pi*_— the PRS of the covariate of interest. For each individual, PRS is constructed by

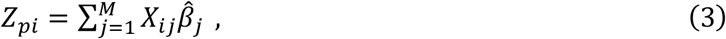

where *M* is the total number of SNPs, *X*_*ij*_ and 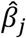 are the genotype and estimated effect size of SNP *j*. We then fit the regression model using *Z*_*p*_ instead to obtain the estimate of the coefficient *γ*. Due to the complexity of genetic architecture and finite sample size of GWAS, *Z*_*pi*_ is often a noisy proxy for *Z*_*i*_.

To allow for the correction of the measurement error, we assume the existence of a small validation dataset where PRS and true value of phenotype are available for *j* = 1, …, *S* individuals. The measurement error can be modeled as:

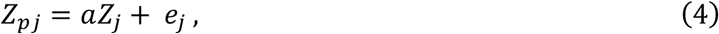

where *a* is the calibration slope of *Y*_*pj*_ regressing on *Y*_*j*_ and *e*_*j*_ ∼ *N*(0, *τ*^5^) is the residual error. The equation above is closely related to the prediction accuracy of PRS. A slope close to 1 indicates PRS is perfectly calibrated and often associated with high prediction accuracy. Since the accuracy of PRS and the calibration slope are influenced by genetic ancestry and many other factors, the validation dataset should have the same ancestry composition as the training dataset used to derive PRS and the testing dataset used to perform the regression.

### Bayesian Approach

Here we describe the full Bayesian model to account for the measurement error of PRS by specifying the distribution *p*(*Z*_*i*_, *γ* | *Z*_*pi*_, *Y*_*i*_) given the observed value of *Z*_*pi*_ and *Y*_*i*_. The joint distribution can be decomposed as

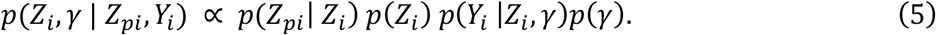

The first component is the measurement error model as formulated below

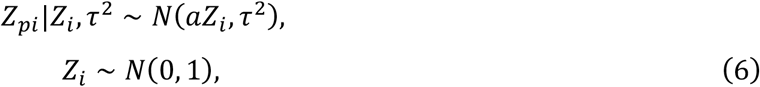

where the slope *a* and the residual variance *τ*^5^ are estimated based on the validation dataset. The second component is the outcome model. For linear regression, we specify the hierarchical model as:

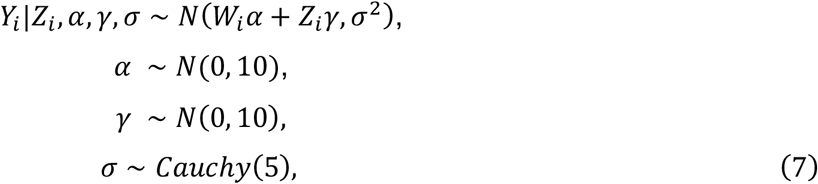

where α and *γ* are regression coefficients. We further assign a non-informative Gaussian prior on α and *γ*. We use a weakly informative normal distribution as the prior for α and *γ*. The prior for parameter σ, representing the standard deviation, is defined by a Cauchy distribution.

Similarly, for logistic regression model, we modeled the binary outcome *Z*_*i*_ in relation to the covariates as below:

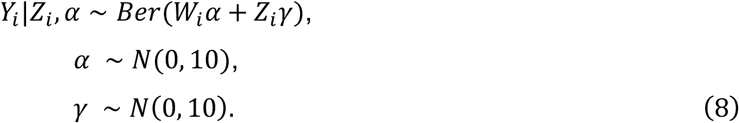

We harnessed Stan, a probabilistic programming language that employs full Bayesian statistical inference through Markov Chain Monte Carlo (MCMC) methods, to fit the above Bayesian model (11). The Stan Model is configured with a total of 2000 iterations per chain. This cumulative 8000-iteration process is preceded by a warm-up phase of 1000 iterations for each chain. This warm-up phase is pivotal in ensuring that the chains achieve a state of convergence, thus establishing a stable foundation for generating meaningful samples for inference. We obtain the posterior mean *E*[*γ* | *Z*_*pj*_, *Y*_*i*_] as the corrected value for the effect size. The credible interval was constructed by picking 2.5% and 97.5% quantile of posterior samples.

### Simulations

In this section, we outline the simulation setup used to generate synthetic data for our analysis. The purpose of simulation is to investigate the performance of our models in recovering inherent relationships among variables and effectively handling measurement errors. We first simulated covariates based on genotypes and derived PRS. We then simulated outcomes based on the linear and logistic regression model.

We used genotypes from the UK Biobank (UKBB) to simulate the covariate of interest (12). A subset of participants from the UKBB were divided into training, validation, and testing datasets each, consisting of 10,000 individuals of European ancestry in each set. We then performed quality control to select 681,828 SNPs for simulation (13). The effect size of each SNP *j* was simulated from a spike and slab prior 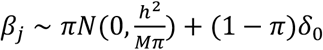, where *h*^2^ = 0.5, *M* = 681,828 and π = 0.01. The covariate of each individual *i* was then generated from the linear model 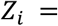 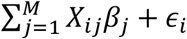 using GCTA-simu with heritability set as 0.5 (14). GWAS summary statistics were then generated for the covariate in the training dataset to construct PRS.

For continuous outcome, we used the linear model to relate *Y*_*i*_ to *Z*_*i*_:

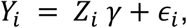

where *γ* is the coefficient of interest, and *ϵ*_*i*_ is the random error following a standard normal distribution.

For binary outcome, we used the logit function to represent the relationship between *Y*_*i*_ and the covariate *Z*_*i*_:

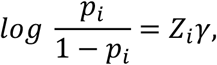

where *p*_*i*_ is the probability of *Y*_*i*_ being 1. *Y*_*i*_ was then sampled from the Bernoulli distribution:

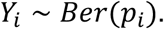

These simulation scenarios provide us with a robust foundation for exploring the performance of our models under different regression settings while considering the presence of measurement errors. For both continuous and binary outcomes, we generated a diverse range of *Y*_*i*_ by varying *γ* from -0.5 to 0.5.

We constructed PRS for the covariate in the testing dataset as 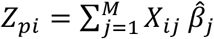, where 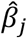 is the effect size of SNP *j* estimated by SDPR based on the generated GWAS summary statistics (13). Subsequently, we employed three strategies to estimate the coefficient *γ* (Figure 1). First, we regressed *Y* on *Z* to obtain the ground truth estimate. Second, we regressed *Y*_*p*_ on Z to obtain the estimate in the presence of measurement error. Third, we applied our Bayesian model to estimate *γ* after accounting for measurement error. We compared the performance of these three strategies in terms of the bias and Bayesian credible intervals of the estimate.

**Figure 1.**
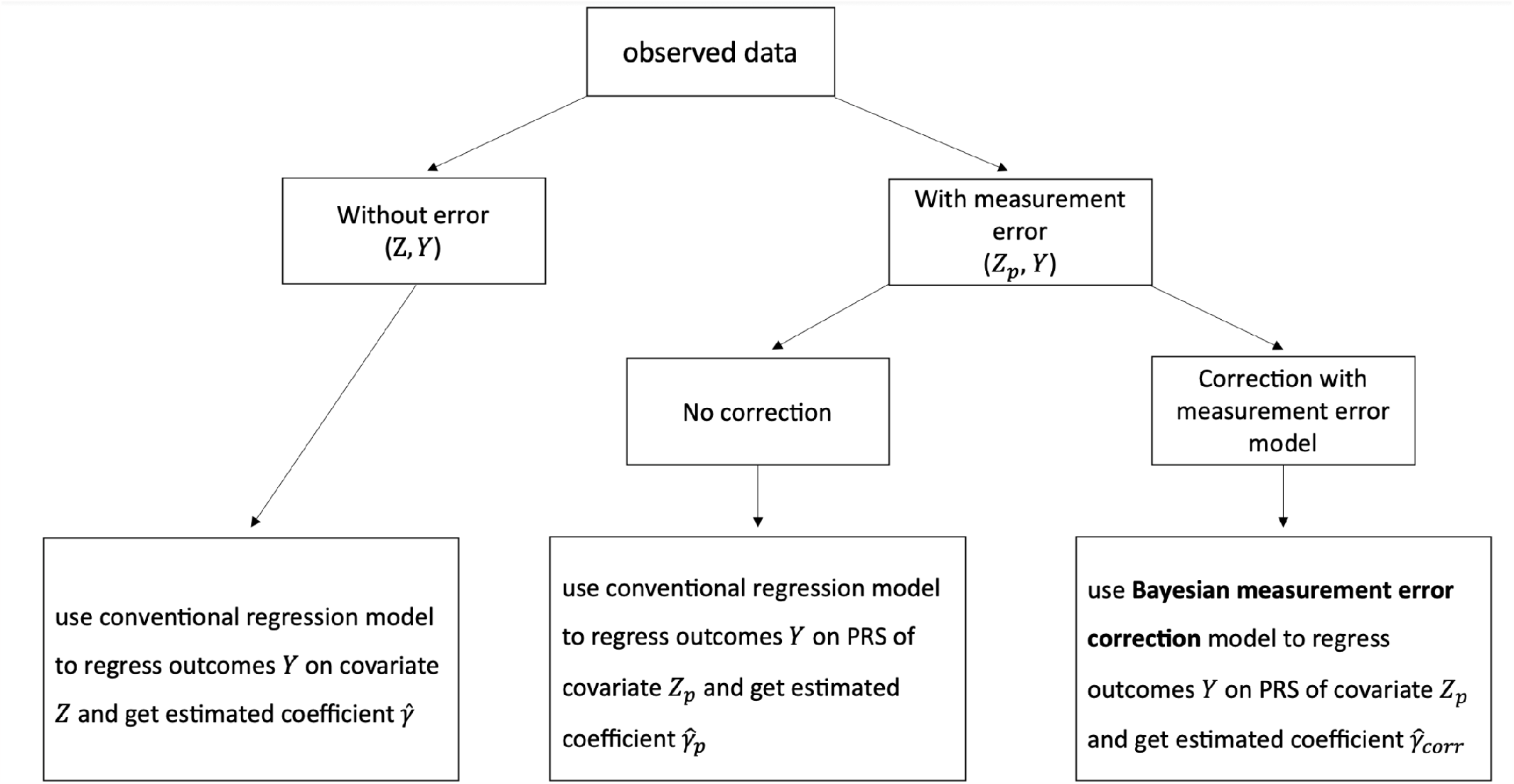
Diagram of three approaches to estimating regression coefficients.

**Figure 2.**
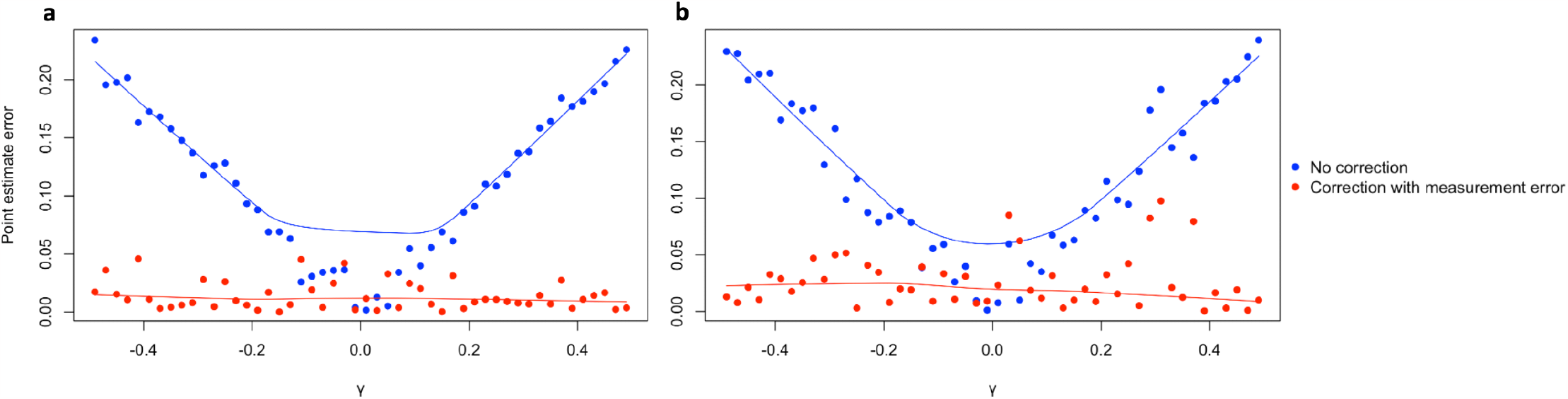
Comparisons of point estimate errors for different simulation scenarios. **a** Continuous outcome simulation results. **b** Binary outcome simulation results. Data are presented as the point estimate error when using the PRS *Y*_*p*_ as the covariate, for the conventional regression models (No correction) and the Bayesian measurement error model (Correction with measurement error) across 50 ground truth *γ* values ranging from -0.5 to 0.5, excluding 0. Both types of outcomes, continuous and binary, are simulated from the models specified in the Simulation section.

### Real Data Applications

We obtained public GWAS summary statistics and trained the PRS model to predict eight traits in the UK biobank (15-20). UK biobank participants with eight traits were selected based on relevant data fields (13). Quality control was performed on summary statistics to remove strand ambiguous (A/T and G/C) SNPs, insertions and deletions (INDELs), SNPs with an effective sample size less than 0.67 times the 90^th^ percentile of sample size. We applied SDPR, a Bayesian nonparametric method that does not rely on specific parametric assumptions on the distribution of effect size, to construct PRS.

For each pair of eight traits, we treated one of them as the covariate and the other one as the outcome. We regressed the outcome on the covariate, adjusting for the additional factor age, to obtain the ground truth estimate of the coefficient. Similar to simulation, we also regressed the outcome on the PRS of the covariate to investigate the impact of measurement error on the estimation. Finally, we applied our Bayesian method to estimate the regression coefficients based on the PRS of the covariate.

## Results

To thoroughly evaluate the efficacy of our Bayesian approach in mitigating measurement error effects within covariates and accurately estimating the inherent relationship between different phenotypes, we conducted comprehensive comparisons of model performance. These assessments involved our Bayesian measurement error model for PRS versus conventional regression models, applied to both synthetic and real-world data in both linear and logistic regression settings.

### Simulation

We first evaluated the performance of the Bayesian measurement error model for PRS on 50 equally sized synthetic datasets. Following the methodology outlined in the Simulation section, we considered both continuous outcome for linear regression models and binary outcome for logistic regression models, by varying the ground truth coefficient *γ* from -0.5 to 0.5, excluding 0 (maintaining equal intervals between consecutive values).

For each dataset, we performed regression of *Y* on *Z*, followed by regressing *Y* on *Z*_*p*_ to obtain coefficient estimates affected by measurement error. As an illustrative example, we set *γ* as 0.1 and the estimated coefficient was 0.106 (95% CI: 0.087-0.125) when regressing *Y* on *Z*.

However, when we fit the model using PRS *Z*_*p*_ as the covariate, we got an estimate of 0.06 (95% CI: 0.036-0.075). This simple example demonstrated that performing regression with PRS as covariate may have downward bias and incorrect CI. Subsequently, we applied the Bayesian measurement error model to regress *Y* on *Z*_*p*_ and derived corrected coefficient estimates and CIs for further comparison.

Table 1 presents the results of coefficient estimations and their corresponding 95% Bayesian credible intervals (CIs) for the three regression models in the linear regression setting. When we employed the Bayesian measurement error model, we observed an 87.9% reduction in the average absolute point estimate error, decreasing from 0.113 to 0.014 compared with regressing *Y* on *Z*_*p*_. At the same time, the probability of 95% CIs covered the true *γ* increased pronouncedly and became the same as the coverage probability for regressing *Z* on *Y*.

**Table 1.** Simulation results for linear and logistic regression. Average point estimate errors and the coverage probability of 95% (Bayesian) credible intervals for three regression scenarios: using the true covariate *Z* (No measurement error), using the PRS *Z*_*p*_ of the covariate (No correction), and using the Bayesian measurement error model to regress on the PRS *Z*_*p*_ of the covariate (Correction with measurement error). The results are based on 50 simulations with a range of ground truth *γ* values ranging from -0.5 to 0.5, excluding 0.

|  | Linear Regression |  |  | Logistic Regression |  |  |
| --- | --- | --- | --- | --- | --- | --- |
|  | No measurement error | No correction | Correction with measurement error | No measurement error | No correction | Correction with measurement error |
| **average point estimate error** | 0.009 | 0.113 | 0.014 | 0.016 | 0.118 | 0.028 |
| **CI containing true $\gamma$** | 94% | 8% | 94% | 98% | 14% | 98% |

Similar to the observations in the linear regression case, the results obtained from logistic regression also showed the efficacy of the Bayesian measurement error model: a 76.5% decrease in the average point estimate error from 0.118 to 0.028 and the 95% CI notably expanded to achieve a 98% coverage of the true *γ* value.

Additionally, as evident from the relatively stable fitted error curve across various *γ* values, when compared to the other fitted curve, whose slope is roughly associated with *γ*, the Bayesian measurement error model successfully resolved the problem of growing absolute error as the magnitude of the ground truth *γ* increased (Figure 1a and Figure1b).

### Real Datasets

#### Linear Regression Analysis

In this section, we first evaluated the performance of the Bayesian measurement error model using datasets for five pairs of continuous traits: Body mass index (BMI), high-density lipoprotein cholesterol (HDL), low-density lipoprotein cholesterol (LDL), total cholesterol (TC), and triglycerides (TG) from UK Biobank. In each of the twelve distinct cases of linear regression analysis, we considered one of the four lipid traits as the outcome variable and utilized one of the remaining three as the covariate variable.

To establish a performance benchmark, we first employed the estimates generated by the built-in linear model in R when regressing the outcome on the covariate as our reference estimator. This reference point allowed us to quantitatively assess the improvements in coefficient estimate accuracy achieved by our approach in each regression case. Subsequently, we conducted a regression of the outcome on the PRS of the covariate with or without using the Bayesian measurement error correction model.

We randomly selected a dataset of size N = 1500 to investigate the linear relationship between height and BMI. In the same manner, we repeated this procedure with a dataset comprising N = 2000 samples to examine the relationships among the four lipid traits (HDL, LDL, TC, and TG).

Supplementary Table 1 and Figure 3(a) present the average point estimate error in coefficient estimates between the conventional linear regression model and the Bayesian measurement error model. The effectiveness of the Bayesian measurement error model in mitigating attenuation bias was evident by the significant decrease in the absolute error for nearly all real trait pairs. However, for one specific trait pair of HDL and LDL, the Bayesian measurement error model did not enhance the estimation accuracy, and using the PRS of the covariate introduced only minimal attenuation bias. This was due to the very small reference coefficient (*γ* = 0.015). For all the other trait pairs, we observed that the Bayesian measurement error model caused a reduction of 48.2% in the absolute error of coefficient estimates on average.

**Figure 3.**
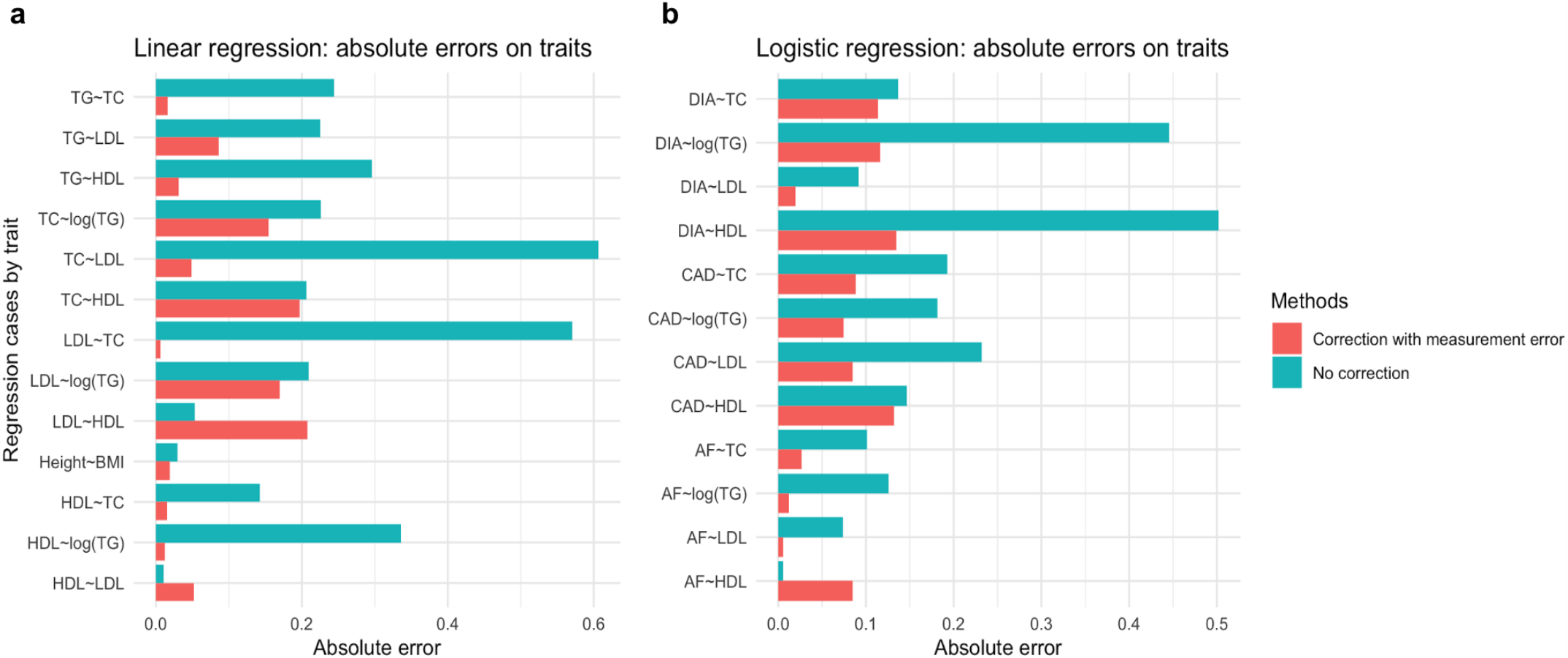
Regression coefficient error comparison among regression cases by traits. **a** Linear regression analysis. **b** Logistic regression analysis. Data are presented as the absolute error in estimated coefficients when using two methods: the conventional regression model (no correction) and the Bayesian measurement error model (correction with measurement error). Errors are computed as the difference between the estimated coefficients when using the PRS and the true value of the covariate.

#### Logistic Regression Analysis

In this section, we evaluated the performance of the Bayesian measurement error model on datasets with binary outcomes. We considered three distinct binary disease outcomes: coronary artery disease (CAD), atrial fibrillation (AF), and type 2 Diabetes (DIA). For each disease outcome, we performed logistic regression on one of the lipid traits as the covariate.

Given that the number of controls is large in UK Biobank, we created a subset of the dataset consisting of all cases and an equal number of controls for more efficient evaluation of different methods. For each dataset, we used both the Bayesian measurement error model and the conventional logistic regression model in Stan to perform regressions of disease outcomes on the PRS of covariates. To obtain the ground truth for model performance evaluation, we utilized the estimates given by the built-in logistic regression model in R by regressing binary outcomes directly on covariates.

Supplementary Table 2 and Figure 3(b) show the results of regressions for all cases, and we observed a significant decrease in absolute errors when using the Bayesian measurement error model. Specifically, across all the eleven regression cases with the underlying true coefficient greater than 0.1, the Bayesian measurement error model achieved a 62.3% reduction in absolute error.

## Discussion

Building on the success of GWAS and the availability of large-scale biobank data, it is increasingly popular to study the relationship across phenotypes using PRS as a surrogate. As we have shown, the use of PRS as the predicted value of the covariate can lead to a downward bias in coefficient estimates for regression models and incorrect construction of the confidence/credible interval. To address this problem, we employed a Bayesian approach to account for the measurement error of PRS and mitigate the attenuation bias in regression models. Our model allows flexible assumptions and can be applied to both continuous and binary outcomes. Through simulations and real data analysis, we showed that our model is effective to reduce the error in coefficient estimates and create the correct credible interval.

Another interpretation for regressing outcome on the PRS of the covariate is to estimate the effect of genetic component of the covariate on the outcome, which has been widely adopted in transcriptome-wide association studies (TWAS) and investigation of gene-environment (G×E) interaction (21-23). In this paper, we did not focus on this interpretation as it would be difficult to establish the ground truth for comparison in the real data analysis. Instead, we treated the coefficient of regressing outcome on the covariate as the ground truth and aimed to recover this true coefficient using PRS as the covariate.

While our method is effective in reducing the attenuation bias, we do note that there are several limitations. First, there is not much improvement for our method when the true regression coefficient is close to 0. Second, the predictive power of PRS cannot be too low, otherwise it is questionable to use it as a covariate in the first place. Third, our method is not able to recover the loss of power due to the noise in PRS. Despite these limitations, our approach has potentially wide applicability in the era of biobank. For example, when the covariate of interest has many missing values, one can first compute the PRS of the covariate, estimate the relationship between PRS and the covariate based on a small dataset, and then apply our approach to correctly obtain the regression coefficient.

## Code availability

Our approach is freely available as an open-source R package available at https://github.com/xinyueq/BayesMEModel.

## Acknowledgements

This work was supported in part by NIH grants R01 HG012735 and R01 GM134005, NSF grant DMS 1902903, and funding from Boehringer Ingelheim. We conducted the research using the UK Biobank resource under an approved data request (ref: 29900). We sincerely thank GIANT, GLGC, CARDIoGRAMplusC4D, BCAC, and DIAGRAM consortia for making their GWAS summary data publicly accessible.

## Supplementary materials

**Supplementary Table 1.** Linear regression coefficients for pairs of continuous traits.

|  | No correction | Correction with measurement error |
| --- | --- | --- |
| HDL~LDL | 0.011 | 0.052 |
| HDL~TC | 0.142 | 0.015 |
| HDL~log(TG) | 0.335 | 0.012 |
| LDL~HDL | 0.053 | 0.208 |
| LDL~TC | 0.571 | 0.006 |
| LDL~log(TG) | 0.209 | 0.170 |
| TC~HDL | 0.206 | 0.197 |
| TC~LDL | 0.606 | 0.049 |
| TC~log(TG) | 0.226 | 0.154 |
| TG~HDL | 0.296 | 0.031 |
| TG~LDL | 0.225 | 0.086 |
| TG~TC | 0.244 | 0.016 |
| Height~BMI | 0.030 | 0.019 |

**Supplementary Table 2.** Logistic regression coefficients for binary outcomes.

|  | No correction | Correction with measurement error |
| --- | --- | --- |
| CAD~HDL | 0.146 | 0.132 |
| CAD~LDL | 0.232 | 0.085 |
| CAD~TC | 0.193 | 0.089 |
| CAD~log(TG) | 0.181 | 0.074 |
| AF~HDL | 0.006 | 0.085 |
| AF~LDL | 0.074 | 0.006 |
| AF~TC | 0.101 | 0.026 |
| AF~log(TG) | 0.126 | 0.012 |
| DIA~HDL | 0.502 | 0.135 |
| DIA~LDL | 0.092 | 0.020 |
| DIA~TC | 0.137 | 0.114 |
| DIA~log(TG) | 0.446 | 0.116 |

